# Biomedical Graph Visualizer for Identifying Drug Candidates

**DOI:** 10.1101/2020.11.27.368811

**Authors:** Ashton Teng, Blanca Villanueva, Derek Jow, Shih-Cheng (Mars) Huang, Samantha N. Piekos, Russ B. Altman

## Abstract

Millions of Americans suffer from illnesses with non-existent or ineffective drug treatment. Identifying plausible drug candidates is a major barrier to drug development due to the large amount of time and resources required; approval can take years when people are suffering now. While computational tools can expedite drug candidate discovery, these tools typically require programming expertise that many biologists lack. Though biomedical databases continue to grow, they have proven difficult to integrate and maintain, and non-programming interfaces for these data sources are scarce and limited in capability. This creates an opportunity for us to present a suite of user-friendly software tools to aid computational discovery of novel treatments through de novo discovery or repurposing. Our tools eliminate the need for researchers to acquire computational expertise by integrating multiple databases and offering an intuitive graphical interface for analyzing these publicly available data. We built a computational knowledge graph focused on biomedical concepts related to drug discovery, designed visualization tools that allow users to explore complex relationships among entities in the graph, and served these tools through a free and user-friendly web interface. We show that users can conduct complex analyses with relative ease and that our knowledge graph and algorithms recover approved repurposed drugs. Our evaluation indicates that our method provides an intuitive, easy, and effective toolkit for discovering drug candidates. We show that our toolkit makes computational analysis for drug development more accessible and efficient and ultimately plays a role in bringing effective treatments to all patients.

Our application is hosted at: https://biomedical-graph-visualizer.wl.r.appspot.com/

## 2. Introduction

The Global Burden of Disease Study conducted in 2013 reported that approximately 95.7% of the world’s population had health problems and 2.3 billion people were experiencing more than five ailments^20^. Even though many of these major health challenges – including cancer, neurodegenerative diseases, and infectious diseases – are in desperate need of drug discovery and innovation, only a third of the around 30,000 currently known diseases have any treatment^17^. Even with the advancement of technology and the accumulation of scientific knowledge, drug discovery still takes an average of 12 to 15 years and costs billions of US dollars^11^.

One of the major bottlenecks for drug discovery is the process of identifying potential drug candidates. Most of the preclinical drug discovery efforts are performed using high throughput screening technologies, which automate the screening of the entire chemical compound library to identify molecules that interact with a particular target of interest^9^. However, high throughput screening does not consider prior knowledge and known pharmacology of the drug candidates, and can only test for a single target^9,12^. It has become increasingly clear in the scientific literature that diseases are driven by multiple molecular abnormalities, and thus identifying drug candidates without prior knowledge is unlikely to be effective^18^. This traditional method of drug discovery requires a tremendous amount of time and resources yet only averages a success rate of 9.6% across all types of diseases^28^.

The exponential growth of biomedical data with heterogeneous data types ranging across several domains including genomics, proteomics, and diseases, has facilitated a new paradigm of drug discovery through computational methods^2^. Computational tools allow researchers to identify and prioritize drug candidates more efficiently, reducing time and resource costs compared with the traditional drug development process^35,15^. Many studies have demonstrated the efficacy of computational methods for drug discovery and repurposing by leveraging publicly available datasets, including: drug focused, disease focused, and ‘omics databases^10,7,13,16,19,5^. However, almost all of these studies used data from a single source, which can introduce biological and technical bias^24^. The multifaceted nature of drug discovery requires intersections of many disparate data types, yet all of the studies mentioned above only used a single type of data.

Though research shows that using multiple data sources can make drug discovery more efficient^14^, integrating these databases and performing algorithmic analysis creates a huge barrier to entry for most drug developers. Integrating and mapping different datasets is a nontrivial task and involves several issues including conflicting nomenclature (brand name vs. generic name) and spelling of medications (American vs. British spellings)^35^. Selecting the appropriate subset from numerous publicly available datasets and creating a data pipeline for drug discovery poses another time-consuming challenge. Glicksberg et al. attempted to mitigate these issues by prescribing a step-by-step workflow for leveraging public data for drug discovery; this framework has proven successful in two separate studies^35,32,31^. However, this method has limited accessibility and utility as it only uses a handful of datasets and requires programming knowledge to execute. Various computational tools have also been made available for linking drug and disease databases, but each has its shortcomings, including: limited data types and sources, limited disease types, and requirements for programming knowledge^21,22,26,29,27,8^. Reducing the number of steps needed for a researcher to begin network analysis on a graph specifically curated for their research task will significantly reduce the barrier to entry for effective computational analysis.

Wikidata is a free and comprehensive knowledge graph covering a variety of concepts and relationships. Researchers have begun to utilize Wikidata for biological reasoning tasks [39, 23, 33]. In this paper we evaluate whether Wikidata is of sufficient quality to construct a biomedical knowledge graph and make inferences about drug use and repurposing. To address the broad problem of access to network analysis, we provide tools that allow users to take advantage of the rich information stored in our biomedical knowledge graph. Our core contribution is to provide two simple, free, and lightweight visual tools for effective network analysis on complex graphs: a subgraph explorer and concept similarity visualizer. We design and extract a biomedical knowledge subgraph from Wikidata and build tools that utilize semantic and clinically salient relationships present in the graph. Our evaluation indicates that our methods can recover known repurposed drugs and that they are intuitive and accessible.

## 3. Data

### 3.1. Wikidata

Wikidata is a free, open, and editable knowledge graph that contains information derived from Wikipedia, third party databases like Uniprot, Gene Ontology, and NCBI, as well as medical literature from PubMed. Wikidata allows programmatic access to its knowledge graph through the use of nodes and edges. A Wikidata node is described by its Wikidata ID, an integer prefixed with a Q (such as Q3025883 for *Type-2 Diabetes*), and contains a list of relationships, or edges, which are described by the property whose identifier is prefixed with a P (such as P31 for *instance of*).

### 3.2. DrugBank

DrugBank is a database comprised of drugs and their known binding proteins [4]. Every drug in DrugBank is a hypothesis for drug candidates that could treat specific medical conditions. DrugBank v5.1.7 contains 13,670 drug entries and 5,228 non-redundant protein (i.e. drug target) sequences linked to these drug entries.

## 4. Methods and Results

### 4.0.1. Extracting Biomedical Data with SPARQL Queries

Because Wikidata stores information in a Resource Description Framework (RDF) triplestore format, the network is stored as semantic triples describing relationships in the form (Concept 1, Property, Concept 2). This allows computational access of its data through SPARQL, an language used to query triple patterns found in graphs. We queried Wikidata using the Python package SPARQLWrapper^40^, and downloaded all instances of biomedical concepts and their relationships enumerated in Figure 2.

**Figure 1.**
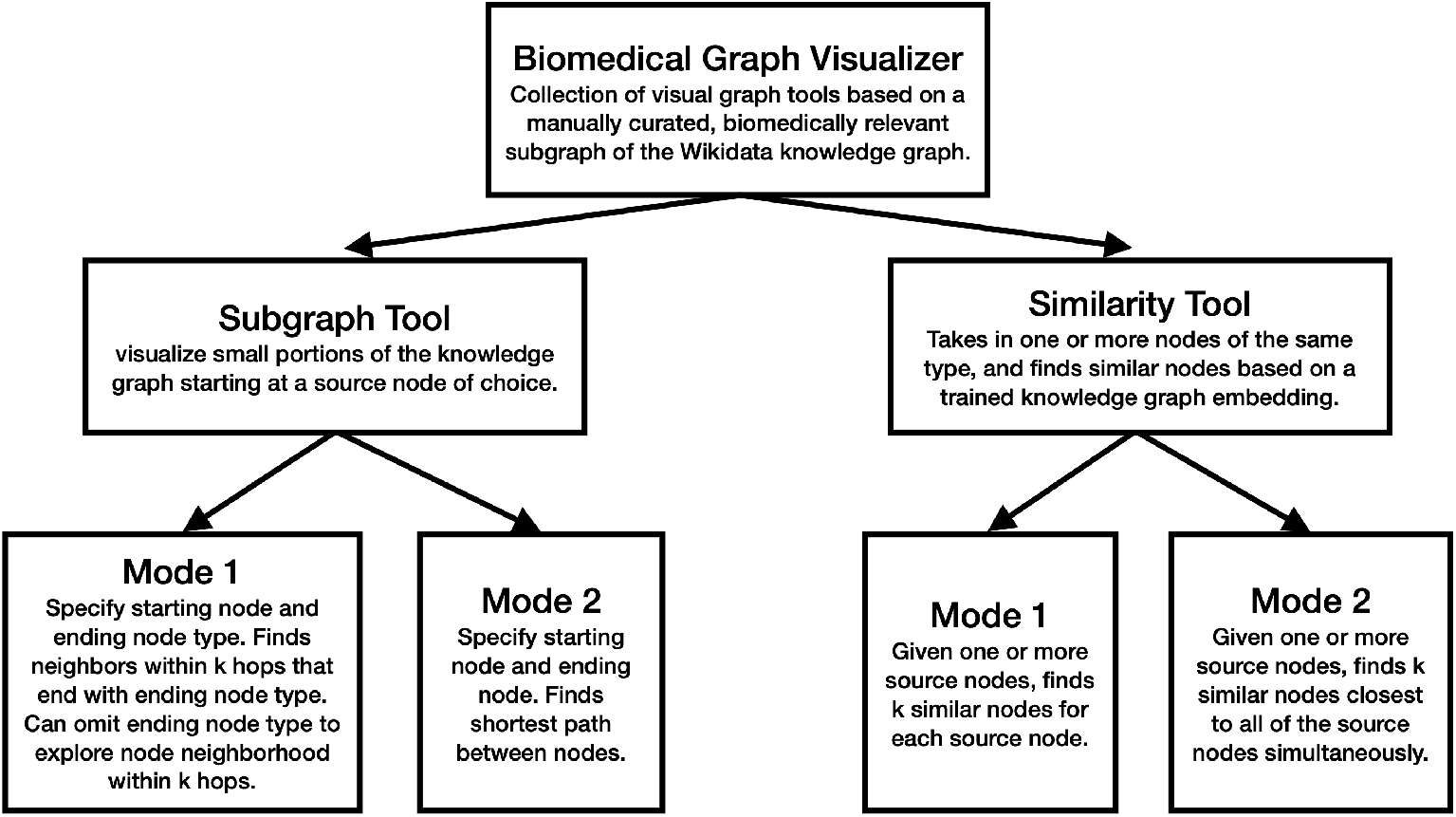
Flowchart of the Biomedical Graph Visualizer, which includes two free visual tools for effective network analysis on complex graphs - the Subgraph Tool and the Similarity Tool. Each also includes 2 modes for performing flexible analyses.

**Figure 2.**
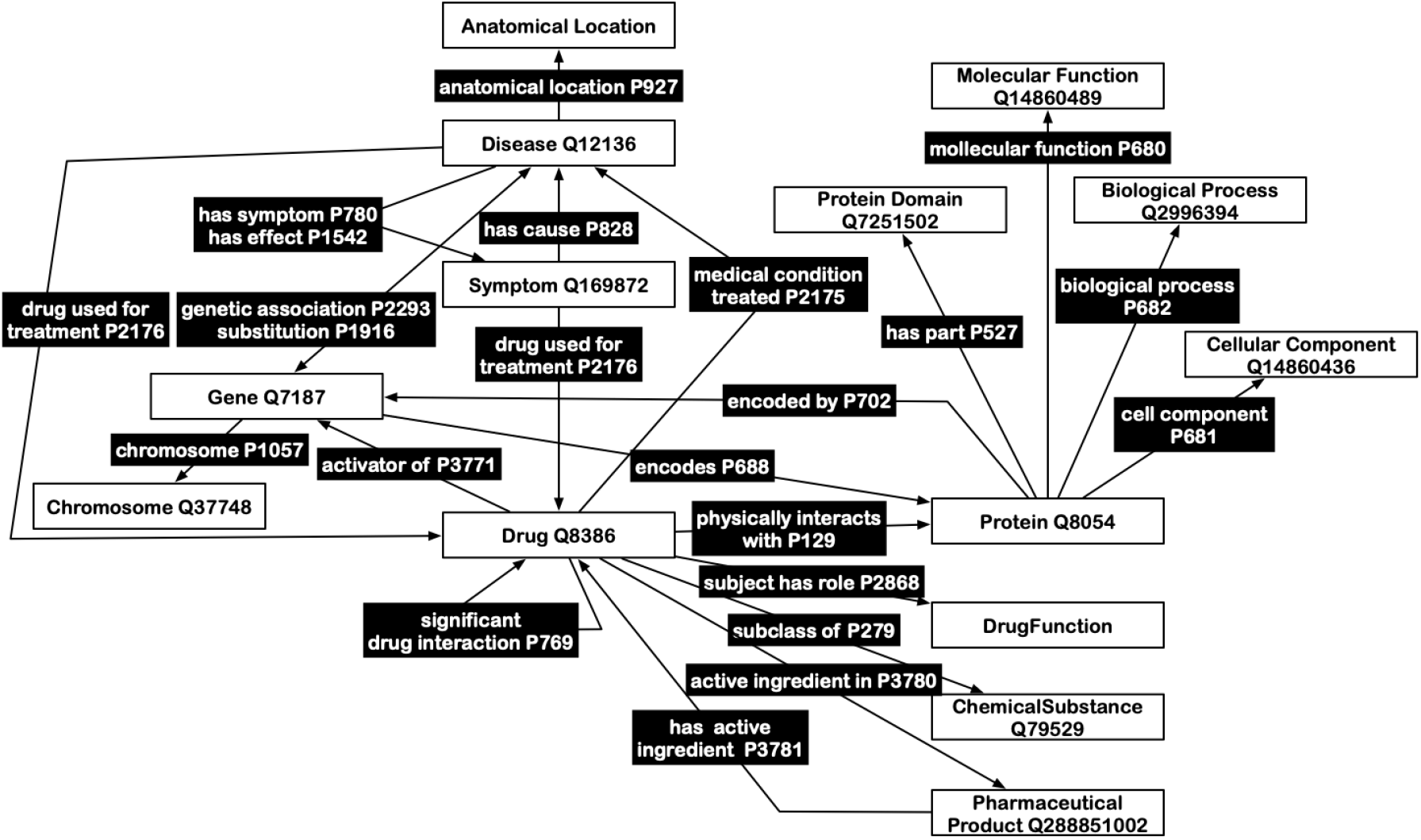
Schema illustrating the concepts and relationships that make up our biomedical knowledge graph. The chosen concepts and relationships were manually curated and taken from Wikidata, and therefore is a subset of Wikidata. Wikidata nodes are in white boxes, and edges display the property ID that connects two nodes together (black boxes). The core concept Drug is centrally located in the subgraph.

Wikidata contains millions of nodes and edges covering a large variety of concepts. We filter the entire Wikidata graph to only include a subset of biomedical concepts and properties useful for drug discovery. The core and peripheral concepts chosen are listed in Table 1. The main distinction between core and peripheral concepts is that all instances of core concepts in Wikidata are included in our graph, whereas peripheral concepts are only included if they have a direct connection via a curated relationship with a core concept.

**Table 1.**
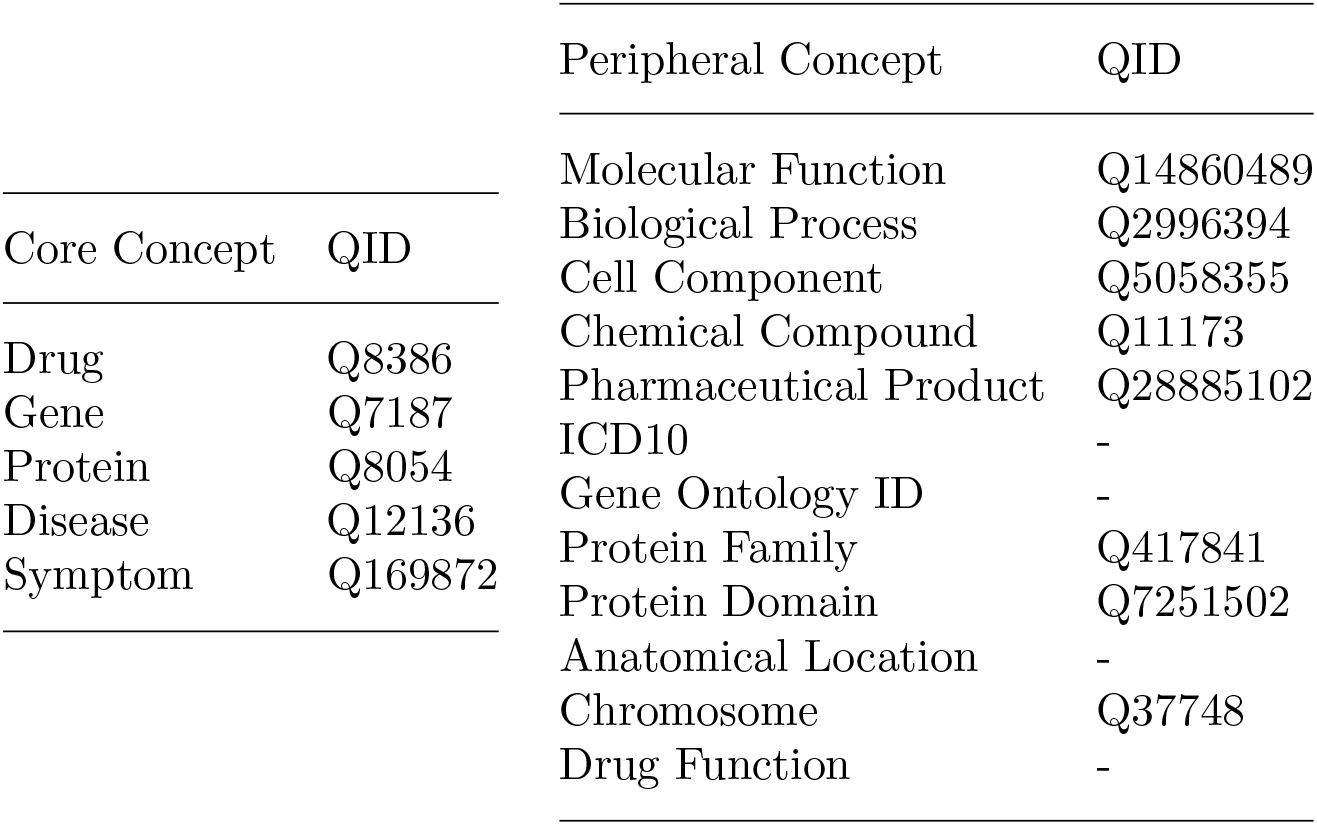
List of core biomedical concepts and peripheral biomedical concepts that we have curated to be a part of our knowledge graph. All instances of core concepts are in our graph, and instances of peripheral concepts are included if they are directly connected to a core concept. Some peripheral concepts do not have a QID.

### 4.1. Biomedical Graph Library

#### 4.1.1. Graph Construction

We use SPARQL queries to download all Wikidata nodes and edges included in our manually-curated schema (the full schema can be seen in Figure 2). SPARQL queries operate on tuples of the form (Concept 1, Property, Concept 2). For example, we can form a SPARQL query to grab all instances of (Gene, encodes, Protein). “Gene” and “Protein” are Concepts, and “encodes” is a Property. In this way, we import all the Concepts listed in table 1, and the Properties that connect them as an adjacency list. Our graph library connects all of edges in this adjacency list to form our biomedical knowledge graph.

We represent this graph as a custom Graph class in Python, based on a NetworkX DiGraph^3^, a directed graph which allows self loops. The DiGraph structure allows us to encode two-sided relationships such as (Gene, encodes, Protein) and (Protein, encoded by, Gene) into one set of bidirectional edges between the same two nodes. An useful feature of NetworkX graphs is that metadata can be added onto each node and edge. We represent each node by its id (e.g. Q17853272) and attach its name (e.g. BRCA2) and concept (e.g. Gene) as metadata on that node. We represent each edge by its id (e.g. P688) and attach its name (e.g. encodes) as metadata on that edge. Furthermore, our Graph class wraps common NetworkX functions such as getting, adding, and deleting nodes and edges. In addition, our library allows getting neighbors, getting metadata, and getting a subgraph, all of which are used by our algorithms. Our custom Graph class abstracts away detailed NetworkX syntax from other parts of our program and preserves the ability to change the underlying graph architecture from NetworkX to other libraries without changing the entire code base.

#### 4.1.2. Graph Analysis and Results

The current version of our biomedical knowledge graph contains 109,230 nodes (concepts) and 570,972 edges (relationships and links between the concepts). See Table 2 for a breakdown of nodes by concept type, and see Figure 3 for a visualization of the distribution of edge types.

**Table 2.** Node counts by concept type in the biomedical knowledge graph. The abundance of nodes in Gene, Protein, Molecular Function and Biological Process suggests that our graph includes an abundance of molecular detail that could describe drug processes.

| Concept | Node Count | Concept | Node Count |
| --- | --- | --- | --- |
| Drug | 2842 | Protein Family | 9017 |
| Gene | 21160 | Protein Domain | 5091 |
| Protein | 25557 | Gene Ontology ID | 11037 |
| Disease | 5074 | Chromosome | 27 |
| Symptom | 604 | Drug Function | 373 |
| Molecular Function | 11095 | Pharmaceutical Product | 2187 |
| Biological Process | 12689 | Cell Component | 1770 |
| ICD10 | 100 | Anatomical Location | 607 |

**Figure 3.**
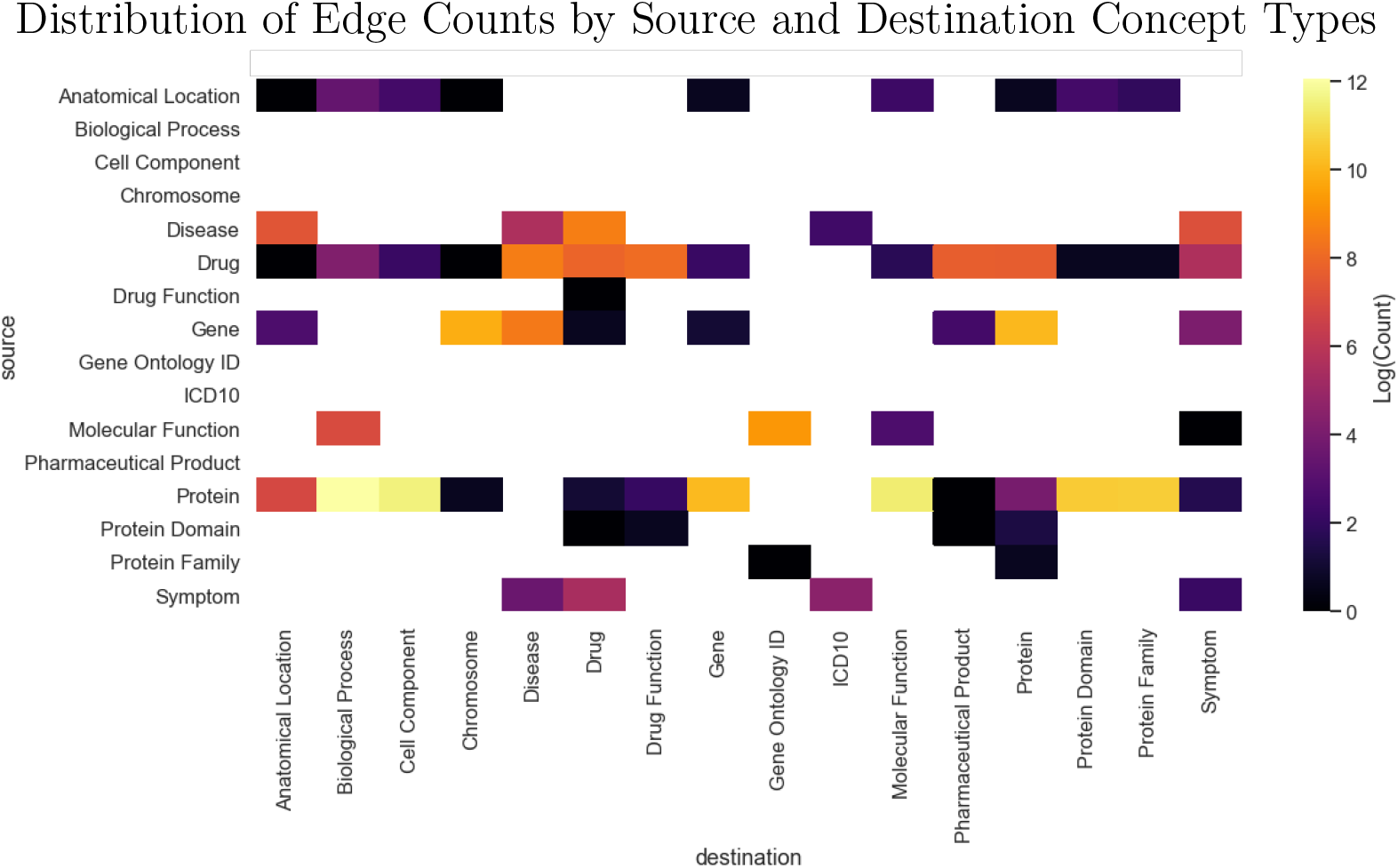
Distribution of edges. Edges in the graph were counted based on the source node concept type and destination node concept type. Natural logarithm was applied to the counts due to the big range, and visualized as a heatmap. White grids denote zero counts in that category. Among the most abundant connections are between Proteins and Biological Processes, Proteins and Molecular Functions, as well as Drugs to Diseases, other Drugs, and Drug Functions. These edges should all be helpful in our drug discovery application.

We investigate the network topology by traversing the graph and recording the distribution of node types encountered at each traversal as in Previde et al. [34]. Figure 4 shows an example of this analysis.

**Figure 4.**
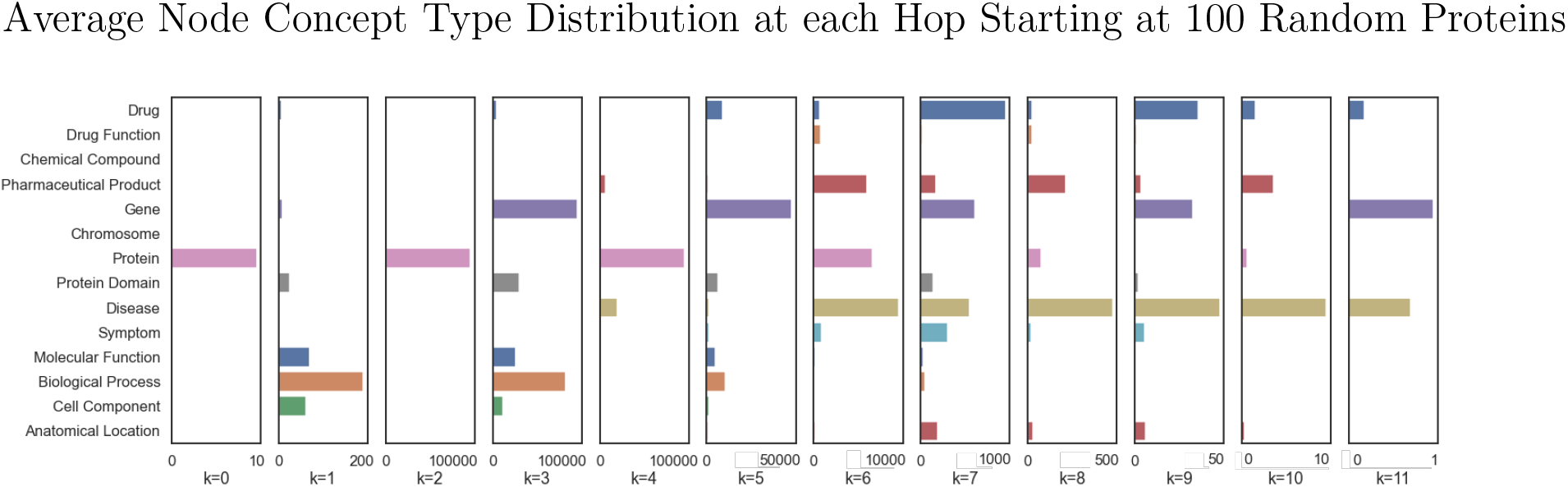
For each of 100 proteins, we conduct BFS and analyze the distribution of neighbor node concept types at each hop of size *k* averaged across the 100 proteins. *k* = 0 refers to the starting protein itself so that for 100 proteins all neighbor node concept types at *k* = 0 should be of type Protein. This analysis shows for proteins, disease concepts are primarily reached after *k* = 6 hops.

### 4.2. Subgraph Tool

#### 4.2.1. Subgraph Computation

The subgraph tool allows users to visualize small portions of the knowledge graph starting at source nodes of their choosing. Users could use it as an exploratory interface for hypothesis generation in drug discovery and repurposing, such as viewing all Drugs that is an activator of a particular Gene of interest. The back-end of the subgraph tool involves three applications of the Breadth First Search (BFS) graph traversal algorithm starting at a specific node that the user specifies. See the Appendix for screenshots of the applications.

The first application explores node neighborhoods, for initial exploration of a concept of interest. The user specifies a source node (e.g. BRCA2) and a maximum number of hops (*k*) to traverse in the graph. The tool will return a subgraph containing the source node and all nodes of all types (e.g. Drugs, Proteins, Diseases) within *k* hops. The interface emulates Previde et al. [34]. BFS is used to explore all adjacent neighbors at each hop before moving to the next set of neighbors. For this application, we keep track of a node’s id and current hop number in the BFS queue. When we pop an item off the queue, we add it to the subgraph and append its neighbors to the queue only if the current hop number is less than *k*.

The second application explores how a specific node is related to a concept domain, suitable for situations when a general relationship between two domains may be hypothesized. The user specifies a source node of interest (e.g. Insulin), an ending concept type (e.g. Disease), and the maximum number of hops (*k*) to traverse. The tool returns a subgraph containing the source node, all nodes of the ending concept type (Disease) within *k* hops, and other nodes on the shortest path between the source node and each ending node. For this application, we keep track of the entire path leading up to the node for each item in the BFS queue, as well as its current hop number. When we pop an item off the queue, we make sure that the current hop number is less than *k*. If the last node in the path is our desired concept, we add all nodes in that path into the subgraph. If not, we continue exploring.

The third application finds the shortest path between two nodes, suitable for finding the relationship between two very specific concepts. Our intuition is that nodes connected by shorter paths are more directly related to each other. We also consider another measure of node relatedness that incorporates relationship types in Section 2.4. The user specifies a source node (e.g. Type-2 Diabetes) and end node (e.g. Liraglutide). The tool returns the nodes and edges on the shortest path between the source and end nodes, irrespective of the number of hops. If the nodes are not connected, the tool will return the source and end nodes with no connections. For this application, we keep track of the entire path leading up to the node for each item in the BFS queue. When we pop an item off the queue, we check if the last node in the path is the ending node specified by the user. If so, we return a subgraph with nodes on this path. Otherwise, we continue exploring.

#### 4.2.2. Subgraph Visualization

Once a subgraph is computed via one of the three methods mentioned above, we employ strategies to visualize the computed subgraphs in an informative way. First, we color the nodes by their concept type and present the users with a labelled legend (not shown). Second, we label the nodes and edges with their names. Both the node and edge labels can be toggled on and off. Third, the size of the nodes represent their estimated connectivity in the graph. For visualization purposes only, we estimate connectivity via PageRank, an algorithm initially developed by Google for ranking web pages on the internet^1^. In our context, PageRank works by counting the number of incoming links at a particular node to determine node importance (centrality). We assume that more important nodes in the graph will receive more connections from other nodes. PageRank assigns a real number between 0 and 1, with higher values denoting higher importance, and therefore a larger node size.

Finally, all of the nodes and edges in the subgraph, along with metadata such as their names, concept types, and PageRank values, are packed into a custom JSON format in an adjacency list form and passed to the web server to be displayed.

#### 4.2.3. Subgraph Tool Evaluation and Results

Since the Subgraph Tool is designed to be intuitive and easily accessible, we evaluated this tool with a user survey. Survey respondents came from biomedical backgrounds. We asked users to pick a biomedical concept and an ending node of their choice. They then rated whether the results were plausible on a scale from 1-5. Survey examples incorporated all three applications of the subgraph tool. Users generally tested queries in fields that were familiar to them. Out of a total of 13 respondents, 5 rated whether the results made sense as 5/5, and 8 rated the results as 4/5. There were no scores lower than 4. Users also wrote open-ended suggestions for what could be improved about the tool. Many users asked for additional features, which are elaborated in the Discussion section.

### 4.3. Similarity Tool

#### 4.3.1. Node Embeddings

The Similarity Tool takes in one or more input nodes of the same type, and generates other similar nodes based on a trained embedding. We generate several node embeddings to compare our results: node2vec, STransE, and attention-based link prediction (henceforth referred to as relation embeddings). Node2vec uses a bag-of-words model and does not take into account relationship types^25^. STransE and relation embeddings take into account edge type (relationship) and direction between nodes^**STransE**, 37^. STransE embeddings are used as a warm-start initialization for the relation embeddings.

We use the default hyperparameters for each embedding algorithm and create embeddings with dim = 50 based on prior work on data with comparable underlying graphs.

Passing our graph to the different embedding algorithms requires minimal processing. We feed an edge list of (src, dst) node tuples to node2vec and an edge list of (src, relation, dst) tuples to the STransE and relation embedding algorithms. For post-processing, we simply split the embeddings by concept type so that only the embeddings required for a particular query are loaded onto memory. The live website uses embeddings produced by the relation algorithm.

#### 4.3.2. Finding Similar Nodes given a Node

The backbone of our similarity tool is the *k* Nearest Neighbors algorithm, which finds nodes similar to each other in our high-dimensional embedding space. Given a source node, the nearest neighbor algorithm returns a list of the *k* most similar nodes based on particular similarity metrics, where *k* is a parameter chosen by the user. For each input source node, distances to all nodes within the same concept (e.g. Drugs) are computed based on our node embeddings. Nodes that have the shortest distance to the input source node are considered to be more similar. The choice of similarity metrics includes (1) cosine distance, (2) Manhattan distance, and (3) Euclidean distance.

Our similarity tool currently supports two different modes: Mode 1 finds a list of nearest neighbors for each of the source nodes independently, while Mode 2 aggregates the results of the individual nearest neighbors and returns a single list of neighbors that are closest to all source nodes simultaneously. The choice of aggregation method includes nearest, mean, and majority. Nearest aggregation concatenates the nearest neighbors for all source nodes, sorts them based on distance, and returns the closest *k* unique neighbors. Mean aggregation takes an average of the distances for a particular node if it appears as a neighbor to multiple source nodes. Majority aggregation sorts the neighbors based on how many times they appear in the top *k* most similar nodes for the source nodes. The variable *k* is user-specified. A user can conduct a similar analysis to that in Figure 4 to determine a reasonable value for *k* for their user case. The value for *k* is capped at *k* = 4 for our graph since higher values of *k* tend to clutter the visualization and hinder readability.

#### 4.3.3. Similarity Tool Evaluation and Results

We evaluate the biomedical graph with a cross-validation and by measuring sensitivity on a held-out set of drug candidate hypotheses from DrugBank. We consider several well-known medical conditions to minimize the effects of noise on the results. The medical conditions we test are: ADHD, Depression, Diabetes, Pulmonary Embolism, Hepatitis B, Heart Attack, and High blood pressure.

##### Cross Validation on the Biomedical Graph

We ran a cross validation on our biomedical graph by taking a set of all nodes 1-hop away from the condition, split the data into *n* = 5 folds (this is equivalent to a 80% train 20% test split of the data), remove all direct edges between the medical condition and drug used for treatment in the test set, generate embeddings for each of the training splits, and evaluate each fold’s STransE embeddings on the respective test set. The results are summarised in Table 3.

**Table 3.** Cross validation to recover known drugs used for treatment of ADHD using only Wikidata as ground truth. We see that our embeddings are able to recover drugs used for treatment of ADHD according to Wikidata. The low mean accuracy (0.13) can be attributed in part to small sample sizes; each fold’s embeddings do not vary significantly from the others as the number of removed edges for each fold is small compared to the overall number of connections in the graph, thus we do not expect performance to vary drastically across runs as the CV graphs are highly similar and folds are not well de-correlated.

| Condition | $n$ | $k$ | Mean acc. | Median acc. | Min acc. | Max acc. |
| --- | --- | --- | --- | --- | --- | --- |
| ADHD | 18 | 5 | 0.13 | 0.00 | 0.00 | 0.33 |

##### DrugBank

We use DrugBank as a test set for evaluating our graph. We measure how well our biomedical subgraph can recover drug candidates listed on DrugBank but not yet present in our biomedical graph as treatment for a chosen set of medical conditions. If the drugs are in Wikidata and DrugBank and have either a “medical condition treated P2175” or “drug used for treatment P2176” edge between Disease and Drug instances in our graph, it is in the train set. If the drugs are in Wikidata and DrugBank and do not have either a “medical condition treated P2175” or “drug used for treatment P2176” edge between condition and drug in our graph, it is in the test set. We take the train and test sets and measure how well our embeddings are. The results are summarised in Table 4.

**Table 4.** Sensitivity results on held-out DrugBank set. We include the highest sensitivity results for each condition using the STransE embedding type, nearest aggregation method, and L2 distance metric, which achieved the highest sensitivity. Many low-sensitivity results had either very few training examples or very few targets, indicating that our test method is susceptible to noise due to insufficient data. With more samples, we expect sensitivity to increase based on the following results.

| Condition | Sensitivity | $n$ |
| --- | --- | --- |
| ADHD | 0.33 | 3 |
| Depression | 0.38 | 21 |
| Diabetes | 0.58 | 33 |
| Heart Attack | 0.06 | 17 |
| Hypertension | 0.22 | 9 |
| Pulmonary Embolism | 0.00 | 4 |

## 5. Discussion

### 5.1. Validation

One main challenge to robust validation is the small set of known drugs used for treatment of medical conditions (for example, our validation set size for ADHD is only *n* = 18); despite this, we are able to successfully recover known drugs used for treatment from both Wikidata and DrugBank data using the embeddings generated from our biomedical knowledge graph. Identifying drugs used for treating medical conditions is a difficult and open problem, and the results discussed in this paper show some success using our approach.

Though our validation methods (recovering known repurposed drugs) were motivated by methods used in prior work^38,36^, larger scale validation is a natural extension of this research. Future work can leverage repurposed drug databases such as repoDB to validate a larger set of conditions than those presented in this paper^30^.

### 5.2. Biomedical Graph Construction

Because Wikidata has millions of nodes, it is likely that we did not include all relevant biomedical concepts. We manually chose gene, protein, drug, disease, and symptom as the core concepts, and then explored several examples of each concept to look at the surrounding node types which we called peripheral concepts. While we believe we captured the majority of biomedical concept nodes related to our application, we do not provide a quantitative analysis of concept coverage and retrieval rates from Wikidata. Similarly, we aimed to capture all the properties that describe how both the core and peripheral concepts are related. We were able to capture the properties through exploring many example nodes, but there are perhaps missing relationships that could be informative in our knowledge graph. Finally, not all property-value pairs are correctly stated in Wikidata. For instance, type-2 diabetes contains “healthy diet” listed under “drug used for treatment”. Therefore, we classify “healthy diet” as a drug. This would require extensive testing to fix this manually for each relationship, so our application will simply display information from Wikidata exactly as is, including any flaws that exist in its underlying structure.

### 5.3. Features and User Interface

As we created a non-programmatic interface to aid users in drug discovery tasks, we necessarily lose the precision and flexibility of programmatic interfaces. In designing features, we had to constantly evaluate the balance between more precise user control over features, and the simplicity and clarity of the website. In our user surveys, many respondents, when performing drug discovery related tasks, asked for more features, such as the ability to export data and interoperability with other systems. For the Subgraph Tool, users asked for ways to block certain relationships in the graph and explore different links in a dynamic way. For the Similarity Tool, users expressed a desire for greater interpretability of the results. Finding the appropriate balance between abstracting away algorithmic details and giving the user sufficient information to make inferences from the displayed results is a design challenge that merits further research.

## 6. Conclusion

We built an intuitive, freely available, and effective toolkit for recovering plausible drug candidates. The survey results and validation discussed above indicate that our knowledge graph, subgraph tool, similarity tool, and web interface jointly make substantial progress towards our goal. Several areas merit further research: User interface; embeddings and nearest neighbor sensitivity; comprehensiveness of the graph; embedding pipeline automation. We hope that further progress on these fronts will provide further evidence that computational methods are central to efficient de novo drug development and repurposing.

### 7. Code Availability

All code related to this project is available at https://github.com/ashtonteng/biomedical-graph-visualizer

## 9. Appendix

### 9.1. User Interface

The website we produced contains the visualizations for our subgraph tool and similarity tool. We used D3.js^6^, a comprehensive Javascript framework for interactive and dynamic visualizations. Images of the pages can be seen in Figures 5, 6, and 7. Our application can be viewed at https://biomedical-graph-visualizer.wl.r.appspot.com.

**Figure 5.**
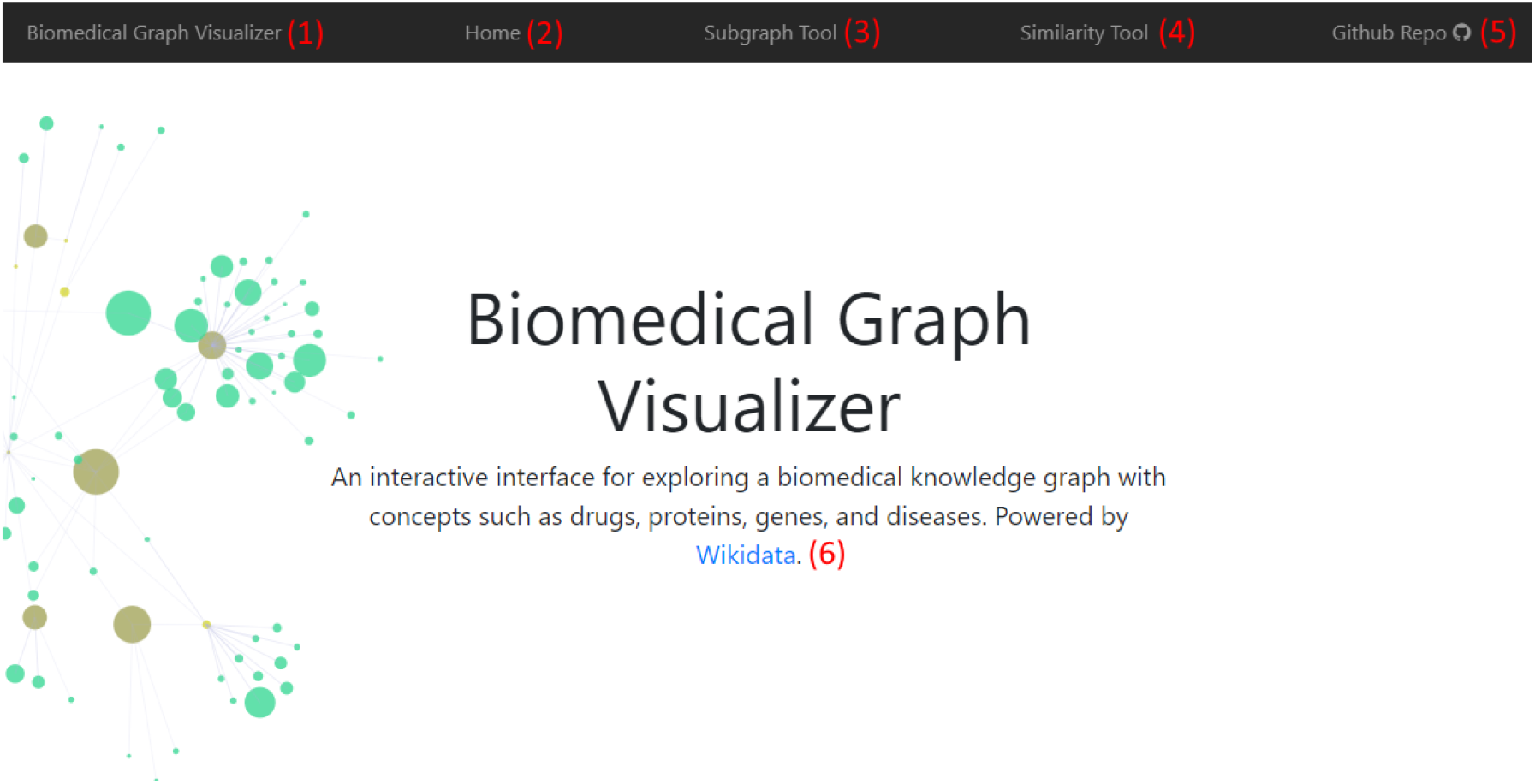
Display of the Home Page for the Biomedical Graph Visualizer tool and tutorial on how to navigate the webpage. The user will see the Bootstrap navigation bar at the top, with each link directing you as follows: (1) Home Page - a portal to all other links and assets used by the tool, including references and how to cite; (2) Home Page; (3) Subgraph Tool - used to visualize subgraphs around biomedical concepts; (4) Similarity Tool - used to visualize similar concepts on a scatterplot; (5) Github Repo - used to inspect and fork the open source code for this project; (6) Wikidata - source of all data used in this project.

**Figure 6.**
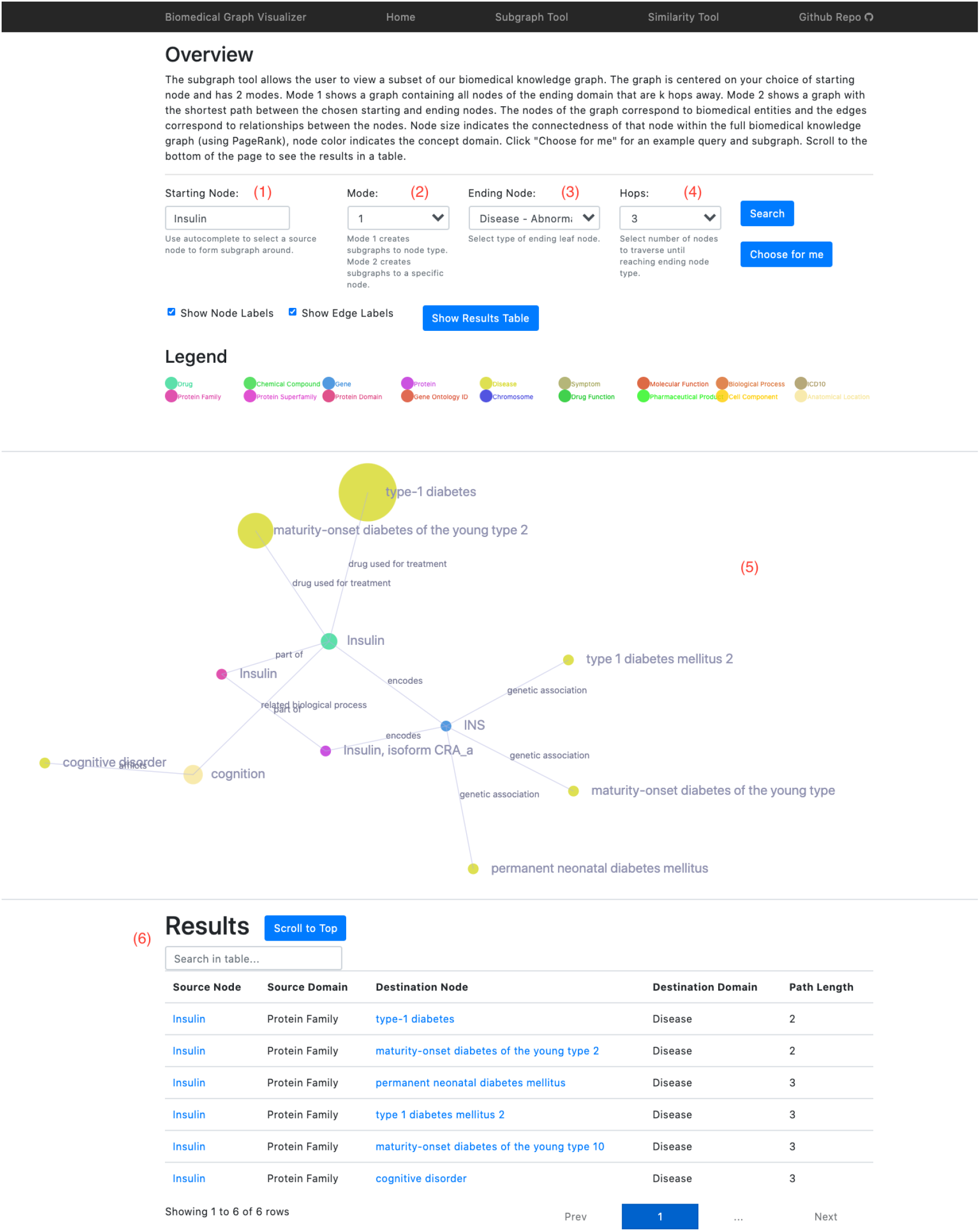
Display of the Subgraph Tool page and tutorial on how to use it. (1) Starting node input form - enter a specific entity here; (2) Mode select - Mode 1 will generate a graph around the concept in (1) to nodes of a given type specified in (3), whereas Mode 2 will generate the shortest path from the entity specified in (1) to the specific entity entered in (3); (3) Ending node input form - takes either a concept type for Mode 1 or specific entity for Mode 2; (4) The depth of the subgraph generated when using Mode 1; (5) D3 force directed graph visualization which will populate a graph according to the parameters specified above when the search button is clicked. Color of each node represents the concept type, whereas the size is proportional to the page rank weight; (6) Table with search bar and pagination, containing an entry for every node rendered in the graph, which will be populated when the search button is clicked.

**Figure 7.**
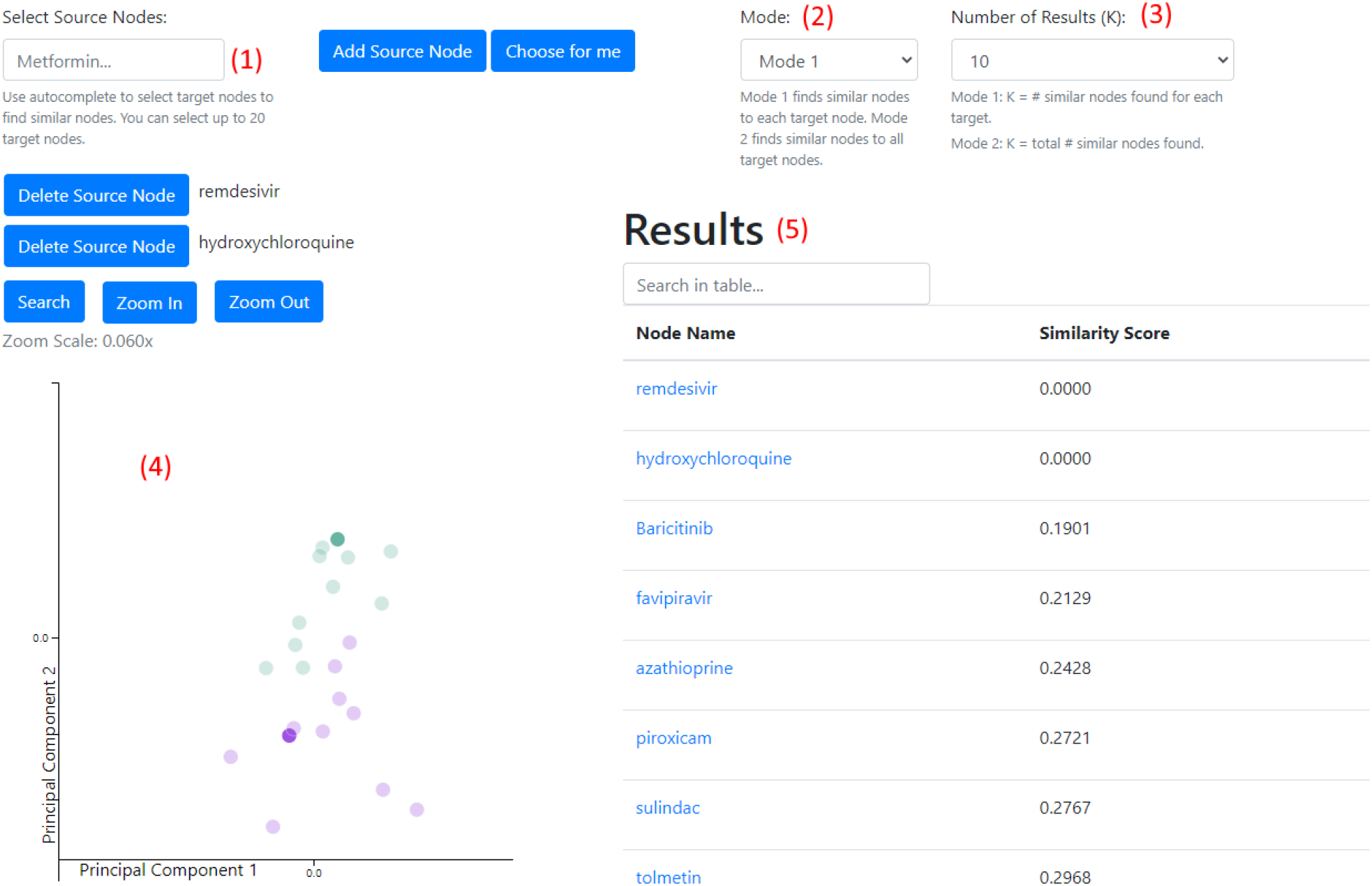
Display of the Similarity Tool page and tutorial on how to use it. (1) Source node form - enter a specific concept here using the autocomplete functionality and click Add Source Node. Repeat this process for each concept to add to the plot, but make sure that the concept types are the same (the colored name that appears in the autocomplete dropdown); (2) Mode selection - Mode 1 computes an individualized KNN on each specified source node and combines the results together (in parallel), whereas Mode 2 will do the same as Mode 1 except the distance is averaged across all specified source nodes; (3) Number of results k; (4) D3 scatterplot visualization which is populated based on the parameters specified above when the search button is clicked. Each source node is assigned a unique color, and the bold intensity points represent the source nodes themselves. In this figure, remdesiver was assigned turqoise whereas hydroxychloroquine was assigned purple - this can be checked by hovering over the point with the cursor. With Mode 1, since a KNN is run for each source node, there will be different classes of points corresponding to each source. In this case, we have two classes, remdesivir and hydroxychloroquine, with turoquoise corresponding to remdesivir and purple corresponding to hydroxychloroquine; (5) Table with search bar and pagination containing an entry for each point on the scatterplot, populated when the search button is clicked. Results are displayed by decreasing distance from the source nodes, indicating that Baricitinib and Favipiravir are similar drug candidates to remdesivir and hydroxychloroquine (Candidate drugs to treat COVID-19).

#### 9.1.1. Front-end Development

The display of the website was created using HTML5, CSS5, and ES6 Javascript. For styling, we used Bootstrap to structure our display. For the subgraph tool, we implemented the D3 force-directed graph, an interactive network widget that allowed us to display the concepts and properties that bind them in a beautiful manner. We were able to specify node color based on node domain and node size based on the output from PageRank^1^. For the similarity tool, we used the D3 scatterplot. We were able to customize point color by index of listed target nodes, and zoom in and out by changing the axis range.

Results from both tools were stored in a dynamic table. We used the publicly available Table Sortable jQuery plugin^41^. This table allowed us to filter results through a search bar at the top and paginate results to display the information in a concise manner.

#### 9.1.2. Back-end Development

We served our templates using Flask^42^, a lightweight Web Server Gateway Interface (WSGI). The Flask server is responsible for serving the HTML, CSS, Javascript, and image files to the user on request.

Additionally, we hosted the web application using Google Cloud Platform. Specifically, we used Google App Engine to run the flask server which dynamically generates a publicly accessible Uniform Resource Locator (URL) to access the application. The virtual machine that is hosting the application is free to use.

## Notes

https://biomedical-graph-visualizer.wl.r.appspot.com

https://github.com/ashtonteng/biomedical-graph-visualizer

### Competing Interest Statement

The authors have declared no competing interest.

## References

[1] Lawrence Page et al. The PageRank Citation Ranking: Bringing Order to the Web. Technical Report 1999-66. Previous number = SIDL-WP-1999-0120. Stanford InfoLab, Nov. 1999. url: http://ilpubs.stanford.edu:8090/422/.

[2] Sandra Kraljevic, Peter J Stambrook, and Kresimir Pavelic. “Accelerating drug discovery: Although the evolution of ‘-omics’ methodologies is still in its infancy, both the pharmaceutical industry and patients could benefit from their implementation in the drug development process”. In: EMBO reports 5.9 (Sept. 2004), pp. 837–842. issn: 1469-221X, 1469-3178. doi: 10.1038/sj.embor.7400236. (Visited on 05/04/2020).

[3] Aric Hagberg, Pieter Swart, and Daniel S Chult. “Exploring network structure, dynamics, and function using networkx”. In: (Jan. 2008).

[4] David S Wishart et al. “DrugBank: a knowledgebase for drugs, drug actions and drug targets.” In: Nucleic acids research 36.Database issue (Jan. 2008), pp. D901–6.

[5] F. Iorio et al. “Discovery of drug mode of action and drug repositioning from transcriptional responses”. In: Proceedings of the National Academy of Sciences 107.33 (Aug. 17, 2010), pp. 14621–14626. issn: 0027-8424, 1091-6490. doi: 10.1073/pnas.1000138107. url: http://www.pnas.org/cgi/doi/10.1073/pnas.1000138107 (visited on 05/04/2020).

[6] Michael Bostock, Vadim Ogievetsky, and Jeffrey Heer. “D^3^ data-driven documents”. In: IEEE transactions on visualization and computer graphics 17.12 (2011), pp. 2301–2309.

[7] J. T. Dudley et al. “Computational Repositioning of the Anticonvulsant Topiramate for Inflammatory Bowel Disease”. In: Science Translational Medicine 3.96 (Aug.x17, 2011), 96ra76– 96ra76. issn: 1946-6234, 1946-6242. doi: 10.1126/scitranslmed.3002648. url: https://stm.sciencemag.org/lookup/doi/10.1126/scitranslmed.3002648 (visited on 05/04/2020).

[8] Assaf Gottlieb et al. “PREDICT: a method for inferring novel drug indications with application to personalized medicine”. In: Molecular Systems Biology 7.1 (Jan. 2011), p. 496. issn: 1744-4292, 1744-4292. doi: 10.1038/msb.2011.26. url: https://onlinelibrary.wiley.com/doi/abs/10.1038/msb.2011.26 (visited on 05/04/2020).

[9] Jp Hughes et al. “Principles of early drug discovery: Principles of early drug discovery”. In: British Journal of Pharmacology 162.6 (Mar. 2011), pp. 1239–1249. issn: 00071188. doi: 10.1111/j.1476-5381.2010.01127. x. url: http://doi.wiley.com/10.1111/j.1476-5381.2010.01127.x (visited on 05/04/2020).

[10] M. Sirota et al. “Discovery and Preclinical Validation of Drug Indications Using Compendia of Public Gene Expression Data”. In: Science Translational Medicine 3.96 (Aug. 17, 2011), 96ra77–96ra77. issn: 1946-6234, 1946-6242. doi:10.1126/scitranslmed.3001318. url: https://stm.sciencemag.org/lookup/doi/10.1126/scitranslmed.3001318 (visited on 05/04/2020).

[11] Peter Csermely et al. “Structure and dynamics of molecular networks: A novel paradigm of drug discovery”. In: Pharmacology & Therapeutics 138.3 (June 2013), pp. 333–408. issn: 01637258. doi: 10.1016/j.pharmthera.2013.01.016. url: https://linkinghub.elsevier.com/retrieve/pii/S0163725813000284 (visited on 05/04/2020).

[12] Jörg Eder, Richard Sedrani, and Christian Wiesmann. “The discovery of first-in-class drugs: origins and evolution”. In: Nature Reviews Drug Discovery 13.8 (Aug. 2014), pp. 577–587. issn: 1474-1776, 1474-1784. doi: 10.1038/nrd4336. url: http://www.nature.com/articles/nrd4336 (visited on 05/04/2020).

[13] V. van Noort et al. “Novel Drug Candidates for the Treatment of Metastatic Colorectal Cancer through Global Inverse Gene-Expression Profiling”. In: Cancer Research 74.20 (Oct. 15, 2014), pp. 5690–5699. issn: 0008-5472, 1538-7445. doi: 10.1158/0008-5472.CAN-13-3540. url: http://cancerres.aacrjournals.org/cgi/doi/10.1158/0008-5472.CAN-13-3540 (visited on 05/04/2020).

[14] Christian Partl et al. “ConTour: Data-Driven Exploration of Multi-Relational Datasets for Drug Discovery”. eng. In: IEEE transactions on visualization and computer graphics 20.12 (Dec. 2014), pp. 1883–1892. issn: 1941-0506. doi: 10.1109/TVCG.2014.2346752. url: https://pubmed.ncbi.nlm.nih.gov/26356902.

[15] JosÃ© P. Pinto et al. “Targeting molecular networks for drug research”. In: Frontiers in Genetics 5 (June 4, 2014). issn: 1664-8021. doi:10.3389/fgene.2014.00160. url: http://journal.frontiersin.org/article/10.3389/fgene.2014.00160/abstract (visited on 05/04/2020).

[16] L. F. Zerbini et al. “Computational Repositioning and Preclinical Validation of Pentamidine for Renal Cell Cancer”. In: Molecular Cancer Therapeutics 13.7 (July 1, 2014), pp. 1929– 1941. issn: 1535-7163, 1538-8514. doi: 10.1158/1535-7163.MCT-13-0750. url: http://mct.aacrjournals.org/cgi/doi/10.1158/1535-7163.MCT-13-0750 (visited on 05/04/2020).

[17] Fischer Birgit. Statistics 2015 - The Pharmaceutical Industry in Germany. Nov. 2015.

[18] Anna Cichonska, Juho Rousu, and Tero Aittokallio. “Identification of drug candidates and repurposing opportunities through compound–target interaction networks”. In: Expert Opinion on Drug Discovery 10.12 (Dec. 2, 2015), pp. 1333–1345. issn: 1746-0441, 1746-045X. doi: 10.1517/17460441.2015.1096926. url: http://www.tandfonline.com/doi/full/10.1517/17460441.2015.1096926 (visited on 05/04/2020).

[19] Hyojung Paik et al. “Repurpose terbutaline sulfate for amyotrophic lateral sclerosis using electronic medical records”. In: Scientific Reports 5.1 (July 2015), p. 8580. issn: 2045-2322. doi: 10.1038/srep08580. url: http://www.nature.com/articles/srep08580 (visited on 05/04/2020).

[20] Theo Vos et al. “Global, regional, and national incidence, prevalence, and years lived with disability for 301 acute and chronic diseases and injuries in 188 countries, 1990–2013: a systematic analysis for the Global Burden of Disease Study 2013”. In: The Lancet 386.9995 (Aug. 2015), pp. 743–800. issn: 01406736. doi: 10.1016/S0140-6736(15)60692-4. url: https://linkinghub.elsevier.com/retrieve/pii/S0140673615606924 (visited on 05/04/2020).

[21] Minjae Yoo et al. “DSigDB: drug signatures database for gene set analysis: Fig. 1.” In: Bioinformatics 31.18 (Sept. 15, 2015), pp. 3069–3071. issn: 1367-4803, 1460-2059. doi: 10.1093/bioinformatics/btv313. url: https://academic.oup.com/bioinformatics/article-lookup/doi/10.1093/bioinformatics/btv313 (visited on 05/04/2020).

[22] Adam S. Brown et al. “ksRepo: a generalized platform for computational drug repositioning”. In: BMC Bioinformatics 17.1 (Dec. 2016), p. 78. issn: 1471-2105. doi: 10.1186/s12859-016-0931-y. url: http://www.biomedcentral.com/1471-2105/17/78 (visited on 05/04/2020).

[23] Sebastian Burgstaller-Muehlbacher et al. “Wikidata as a semantic framework for the Gene Wiki initiative”. In: Database 2016 (2016).

[24] B Chen and Aj Butte. “Leveraging big data to transform target selection and drug discovery”. In: Clinical Pharmacology & Therapeutics 99.3 (Mar. 2016), pp. 285–297. issn: 00099236. doi: 10.1002/cpt.318. url: http://doi.wiley.com/10.1002/cpt.318 (visited on 05/04/2020).

[25] Aditya Grover and Jure Leskovec. node2vec: Scalable Feature Learning for Networks. 2016. arXiv: 1607.00653 [cs.SI].

[26] Riku Louhimo et al. “Data integration to prioritize drugs using genomics and curated data”. In: BioData Mining 9.1 (Dec. 2016), p. 21. issn: 1756-0381. doi: 10.1186/s13040-016-0097-1. url: http://biodatamining.biomedcentral.com/articles/10.1186/s13040-016-0097-1 (visited on 05/04/2020).

[27] Soheil Moosavinasab et al. “‘RE:fine drugs’: an interactive dashboard to access drug repurposing opportunities”. In: Database 2016 (2016), baw083. issn: 1758-0463. doi: 10.1093/database/baw083. url: https://academic.oup.com/database/article-lookup/doi/10.1093/database/baw083 (visited on 05/04/2020).

[28] Asher Mullard. “Parsing clinical success rates”. In: Nature Reviews Drug Discovery 15.7 (July 2016), pp. 447–447. issn: 1474-1776, 1474-1784. doi: 10.1038/nrd.2016.136. url: http://www.nature.com/articles/nrd.2016.136 (visited on 05/04/2020).

[29] Giorgio Valentini et al. “RANKS : a flexible tool for node label ranking and classification in biological networks”. In: Bioinformatics 32.18 (Sept. 15, 2016), pp. 2872–2874. issn: 1367-4803, 1460-2059. doi: 10.1093/bioinformatics/btw235. url: https://academic.oup.com/bioinformatics/article-lookup/doi/10.1093/bioinformatics/btw235 (visited on 05/04/2020).

[30] Adam S Brown and Chirag J Patel. “A standard database for drug repositioning”. In: Scientific data 4.1 (2017), pp. 1–7.

[31] Bin Chen et al. “Computational Discovery of Niclosamide Ethanolamine, a Repurposed Drug Candidate That Reduces Growth of Hepatocellular Carcinoma Cells In Vitro and in Mice by Inhibiting Cell Division Cycle 37 Signaling”. In: Gastroenterology 152.8 (June 2017), pp. 2022– 2036. issn: 00165085. doi: 10.1053/j.gastro.2017.02.039. url: https://linkinghub.elsevier.com/retrieve/pii/S0016508517302640 (visited on 05/04/2020).

[32] Li Li et al. “Novel Therapeutics Identification for Fibrosis in Renal Allograft Using Integrative Informatics Approach”. In: Scientific Reports 7.1 (Apr. 2017), p. 39487. issn: 2045-2322. doi: 10.1038/srep39487. url: http://www.nature.com/articles/srep39487 (visited on 05/04/2020).

[33] Tim E Putman et al. “WikiGenomes: an open web application for community consumption and curation of gene annotation data in Wikidata”. In: Database 2017 (2017).

[34] P. Previde et al. “GeneDive: A gene interaction search and visualization tool to facilitate precision medicine”. In: Pac Symp Biocomput 23 (2018), pp. 590–601.

[35] Benjamin Glicksberg et al. “Leveraging Big Data to Transform Drug Discovery”. In: vol. 1939. Jan. 2019, pp. 91–118. isbn: 978-1-4939-9088-7. doi: 10.1007/978-1-4939-9089-4_6.

[36] Tareq B. Malas et al. “Drug prioritization using the semantic properties of a knowledge graph”. In: Scientific Reports 9.1 (Apr. 2019), p. 6281. issn: 2045-2322. doi: 10.1038/s41598-019-42806-6. url: https://doi.org/10.1038/s41598-019-42806-6.

[37] Deepak Nathani et al. “Learning Attention-based Embeddings for Relation Prediction in Knowledge Graphs”. In: Proceedings of the 57th Annual Meeting of the Association for Computational Linguistics. Florence, Italy: Association for Computational Linguistics, 2019.

[38] K. Park. “A review of computational drug repurposing”. In: Transl Clin Pharmacol 27.2 (June 2019), pp. 59–63.

[39] Andra Waagmeester et al. “Science Forum: Wikidata as a knowledge graph for the life sciences”. In: ELife 9 (2020), e52614.

[40] et. al. Alonso. SPARQLWrapper. url: https://github.com/RDFLib/sparqlwrapper.

[41] Paginate, Filter, And Sort Dynamic Data In A Table - Table Sortable. Version V2.1. JQUERY PLUGINS. url: https://www.jqueryscript.net/table/Paginate-Sort-Filter-Table-Sortable.html.

[42] Armin Ronacher. Flask. Version 1.1.x. url: https://flask.palletsprojects.com/en/1.1.x/#.

